# Computational analysis of the effect of a binding protein (RbpA) on the dynamics of *Mycobacterium tuberculosis* RNA polymerase assembly

**DOI:** 10.1101/2024.02.19.580928

**Authors:** Sneha Bheemireddy, Ramanathan Sowdhamini, Narayanaswamy Srinivasan

## Abstract

RNA polymerase-binding protein A (RbpA) is an actinomycetes-specific protein crucial for the growth and survival of the pathogen *Mycobacterium tuberculosis*. Its role is essential and influences the transcription and antibiotic responses. However, the regulatory mechanisms underlying RbpA-mediated transcription remain unknown. In this study, we employed various computational techniques to investigate the role of RbpA in the formation and dynamics of the RNA polymerase complex.

Our analysis reveals significant structural rearrangements in RNA polymerase happen upon interaction with RbpA. Hotspot residues, crucial amino acids in the RbpA-mediated transcriptional regulation, were identified through our examination. The study elucidates the dynamic behavior within the complex, providing insights into the flexibility and functional dynamics of the RbpA-RNA polymerase interaction. Notably, potential allosteric mechanisms, involving the interface of subunits *α*1 and *α*2 were uncovered, shedding light on how RbpA modulates transcriptional activity.

Finally, potential ligands meant for the *α*1-*α*2 binding site were identified through virtual screening. The outcomes of our computational study serve as a foundation for experimental investigations into inhibitors targeting the RbpA-regulated dynamics in RNA polymerase. Overall, this research contributes valuable information for understanding the intricate regulatory networks of RbpA in the context of transcription and suggests potential avenues for the development of RbpA-targeted therapeutics.

**Author Summary:** Infection studies by *Mycobacterium tuberculosis* (Mtb) acquires primary importance due to its severe infection and antibiotic resistance. There is an open need for highly effective drugs and one needs to employ novel approaches such as detailed structural analysis and the possibility to focus on allosteric inhibitors. We have exploited the availability of cryo-EM structures of RNA polymerase of Mtb, with and without its transcription-activator protein namely RNA polymerase-binding protein A (RbpA). In this study, we employed various computational techniques to investigate the role of RbpA in the formation and dynamics of the RNA polymerase complex. The assemblies were subject to molecular dynamics and perturbation scanning, followed by structural comparisons and measurement of subunit interface strength. These analyses could clearly show that α subunits, which are far away from the RbpA binding site, undergo differential structural changes. Hence, we focused on the αα’ site to recognize potential small molecule inhibitors using virtual screening. These analyses demonstrate that it is possible to perform comparative structural analysis of different forms of assemblies, which can be useful towards drug design.

## 1 INTRODUCTION

Prokaryotes are single-celled organisms and lack distinct nuclei or organelles, yet they manage to perform all essential functions using intracellular organization [1,2]. They depend on large macromolecular complexes for all critical processes ranging from cell division to differentiation [3]. In general, macromolecules do not function in isolation but form functional complexes in association with other macromolecules. The disruption of these macromolecular complex interaction networks forms the molecular basis for many diseases [4]. Hence it is imperative to study their properties such as complex formation, kinetics, and dynamics [5]. It is required to determine its 3-dimensional structure for a detailed characterization of macromolecular biochemical and biophysical behavior. Biophysical techniques like NMR, X-ray crystallography, and cryo-EM [6] can unearth the structural intricacies of the proteins, but every method has its limitations even though techniques in this area have undergone many advancements during the past few decades [7]. Many membrane protein complexes are difficult to crystallize [8], NMR studies are limited by the molecular weight of the complex, and cryo-EM is challenging when protein size is small (<150 kDa). In this manuscript, we discussed the computational analysis of one such macromolecular assembly, namely a bacterial polymerase.

Infection by *Mycobacterium tuberculosis* (*Mtb*) leads to 10 million cases of tuberculosis (TB) annually (Global Tuberculosis Report, 2013). The emergence of multidrug-resistant (MDR) strains presents an even more deadly threat, given the low success rate of the existing antibiotics. One of the key antibiotics used in TB treatment is rifampicin (Rif), a broad-spectrum antibiotic that inhibits the enzyme RNA polymerase (RNAP). It is suggested that resistance to rifampicin leads to more treatment failures and patient deaths than resistance to any other agent (Global Tuberculosis Report 2013, 2013). Actinobacteria have unique transcription factors, including CarD [9,10,11] and RbpA [12,13,14] which cooperatively stabilize the RPo, and as a result, have been characterized as transcriptional activators. RbpA is upregulated when subjected to bacterial stresses and is essential for growth under nutrient-rich conditions [13]. RbpA is vital for maintaining cellular homeostasis and modulating *Mtb* gene expression to drive the pathogenesis of tuberculosis. As RNAP is a direct target for first-line antibiotics used to treat *Mtb* infection, further mechanistic studies of mycobacterial transcription may lead to novel therapeutic strategies.

The specificity of RbpA, the activator, towards sigma factors, has led to theories that RbpA might play a role in the competition for sigma factors to interact with the RNAP core [13]. RbpA was proposed to act in part, by anchoring promoter DNA to RNA Polymerase. Furthermore, RbpA can increase the affinity between the holoenzyme and the promoter, *via* the interaction of the BL with DNA. An arginine residue (R79) in the BL is thought to play a role in RbpA’s ability to bind DNA, although it is unknown whether this interaction may contribute to RbpA’s promoter specificity [14]. *In vivo* studies in *Mycobacterium smegmatis* (*Msm*) confirm that RbpA is essential, and the deletion of constituent domains causes small to severe growth defects. In contrast to *Msm*, *α* R79A mutant in *Mtb* is not viable [12]. The role of RbpA as an activator is already established. It was suggested that the N-terminal tail (NTT) and C-terminal Domain (CD) inhibit RPo formation yet deleting both these domains led to a global decrease in gene expression. Thus, RbpA appears to have pleiotropic functions in transcription and with its mechanism of action unknown.

By exploiting the availability of structures of the polymerase in the bound and unbound forms, and gaining insight into the functional implications of RbpA, we performed a computational study to investigate various aspects of the RbpA-RNA polymerase complex. Molecular dynamics simulations and analysis techniques were employed to elucidate the structural changes occurring within the RNA polymerase complex upon RbpA binding. The analysis revealed significant structural rearrangements in the RNA polymerase complex upon interaction with RbpA. These changes provided insights into the conformational dynamics and stability of the complex. Furthermore, the inter-subunit interactions were found to be altered, indicating the influence of RbpA on the overall architecture and assembly of the complex. Examination of hotspot residues highlighted key amino acids involved in the RbpA-mediated regulation of transcription. Fluctuation analysis demonstrated dynamic behavior within the complex, shedding light on the flexibility and functional dynamics of the RbpA-RNA polymerase interaction. Additionally, the investigation of allosteric effects revealed potential mechanisms through which RbpA modulates transcriptional activity. Additionally, the investigation of allosteric effects revealed potential mechanisms that include the interface of subunits of *α*1 and *α*2 through which RbpA modulates transcriptional activity. We identified ligand binding sites on the interface of subunits of α1 and *α*2. We performed molecular docking using FDA-approved ligands and natural ligands from Molport. We could identify ligands that interact with *α*1 at the binding site of *α*2. The results of this computational study can provide a starting point for experimental investigations on inhibitors of RbpA-regulated dynamics in RNA polymerase.

## 2 MATERIALS AND METHODS

### 2.1 Dataset preparation

We have identified two structures corresponding to *Mtb* polymerase in PDB: RbpA bound holoenzyme (PDB code: 6C04 [15] and RbpA unbound holoenzyme (PDB code: 5UH5 [16] complex. A few residues (1011-1021) of the *β*’ chain was found missing in the PDB files. Hence, we used loop modeling from MODELLER [17] to address this. A bunch of 100 models were generated and a model with the highest DOPE score was selected for further analysis. For molecular dynamics studies, we have made two structures from these structures by removing a subunit *in-silico*. We also considered RNA polymerase bound with RbpA and carD (PDB code: 6EDT) for MD studies.

### 2.2 Structural features

RMSD (Root mean square deviation) was calculated both at the global and individual C*_α_* levels. Global RMSD was calculated by taking the mean RMSD among all the residues and the C*_α_* deviation was measured at residue level. We employed TM-align [18] for both the global and C*_α_* RMSD calculations. If any two atoms (from two different proteins) fall within a distance cut-off (4.5Å or 5Å), residues corresponding to these atoms were regarded as interfacial residues. We used PIC [19] an in-house web server, to identify the interfacial residues using both distance (5Å) and solvent accessibility criteria (less than 7% solvent accessibility). Here we used PPcheck [20] for the calculation of interaction energy. Residues that resulted in energy change greater than 2 kcal/mol upon mutation to alanine were considered hotspot residues [21]. We used PPcheck [20] for the identification of these hotspot residues.

### 2.3 Molecular dynamics analysis

Six different systems were considered for the molecular dynamics (MD) simulations in this study. The starting complex was the RNA polymerase holoenzyme (PDB ID: 5UH5). From this complex, two additional systems were generated: one comprising the core enzyme after removing the sigma factor (*σ*), and another consisting of the RNA polymerase holoenzyme without DNA. Furthermore, a complex without DNA was derived from the RbpA-bound holoenzyme structure (PDB ID: 6C04). Additionally, the RbpA and CarD-bound holoenzyme complex (PDB ID: 6EDT [22] was also included in the analysis. All complexes were carefully prepared by modeling missing residues, ensuring that they possessed an equal number of residues. The DNA present in the complexes was obtained from the 6EDT structure, as it provided a complete open DNA conformation model within the structures. The analysis of MD trajectories involved various parameters and calculations using the MDAnalysis package [23,24].

### 2.4 Perturbation response scanning (PRS)

PRS is based on applying this Linear Response Theory (LRT) to understand the basis for structural changes in proteins. Proteins were represented as an elastic network model using C_α_ atoms. A force of magnitude ΔF unit was applied on the C_α_ atom of each residue, and the direction of the force applied was chosen randomly (isotropic). The whole protein is scanned for the response of a single perturbation. The result of perturbation was recorded considering both the magnitude and direction of the response. These perturbations are repeated at least 1000 times on each residue, and the average of all responses is considered for further analysis. In this study, we have used the ProDy package [25] for PRS analysis of MD data.

### 2.5 Virtual screening

The structures of α subunits obtained from the RbpA-bound RNA polymerase complex were used for the docking studies. Molprot (database of natural products) [26] and FDA-approved molecules dataset [27] were used for ligand screening. Since no information was available about the binding sites, the interface of α subunits was considered, and the binding sites were predicted using Schrodinger’s sitemap tool [28,29]. The docking was performed using Glide [30,31,32]. Initially, HTVS (High Throughput Virtual Screening) protocol was used followed by SP (Standard Precision) docking of top ligands from HTVS based on docking score followed by XP (extra precision) docking of top scoring ligands from SP docking. *ð*G was calculated using the prime MM-GBSA module of Schrodinger. A grid box covering the entire interface was constructed to define the docking space. The number of GA runs was set to 100 in the docking parameters and the Lamarckian genetic algorithm was used [33]. For each conformer, 100 poses of the top 10 ligands were generated.

All the docked structures were simulated in an aqueous environment. The simulation was run using the Desmond package of Schrodinger [34]. The structures were initially prepared using the protein preparation wizard of Schrodinger which involves protonation, H-bond optimization, and restrained minimization [35]. In the system builder, the TIP4P water model was specified for the solvation of the system. The system was neutralized and an ionic concentration of 0.15 mM using NaCl was specified in the system builder. The output of the system builder was relaxed using the default relaxation protocol followed by production MD run for 50 nanoseconds using the OPLS3e force field [36]. For all the structures the MD simulation was run for 100 ns. The simulation interaction diagram (SID) module of the Desmond package was used to analyze the simulation results. The entire range of simulation time was considered for all analyses. RMSD is calculated for each frame by aligning the complex to the protein backbone of the reference frame which is 10^th^ ns. For ligand RMSD, the protein-ligand complex is aligned to the reference frame and then the RMSD of ligand heavy atom is calculated. Significantly higher values of ligand-RMSD than protein- RMSD signifies the diffusion of ligand away from its initial binding site. Protein-ligand interactions were also monitored throughout the simulation time. These interactions include H-bond, Hydrophobic interaction, ionic interaction, and water bridges. Hydrophobic interaction also includes *л*-Cation and *л* − *л* interactions. Custom Python scripts were written to plot these interactions.

## 3 RESULTS AND DISCUSSION

### 3.1 Interaction of the RbpA influences the conformation of the constituents of RNA polymerase complex

To investigate the structural differences in the various RNA polymerase subunits upon interaction with RbpA, tools such as TM- align, DALI [37] and PDBefold [38] were employed. All the methods indicated similar changes in RMSD. So, we went ahead with the standard TM-align for Global RMSD calculations and interface comparison. We observed that the global RMSD ranges from 0.7Å to 2Å (Figure 1), with the former corresponding to subunit *ω* and the latter corresponding to the *σ*-factor of RNA polymerase. We have also observed significant structural changes in *β*’. Notably, RbpA directly interacts with the *σ*-factor and *β*’. These results suggest that RbpA binding induces structural alterations in the subunits directly interacting within the complex. Analysis of the local structural deviations implies that there are more significant structural changes in the local regions of the subunits. Furthermore, through the analysis of local C*_α_* RMSD (as depicted in Supplementary figure S1), the identified residues display notable local structural variations. We conducted a comprehensive analysis of interfacial residues and the interaction energy between different subunits using PPcheck. Our investigation revealed a remarkable reduction in the number of interfacial residues between *β* and *σ* (Figure 1 A), as well as between *β*’ and *σ* upon RbpA binding. Conversely, there was an increase in the number of interfacial residues between *α*1-*α*2 and *α*2 - *β*, upon interaction with RbpA. Although further investigations are required, these observations potentially imply the occurrence of allosteric effects within the RNA polymerase complex.

**FIGURE 1.**
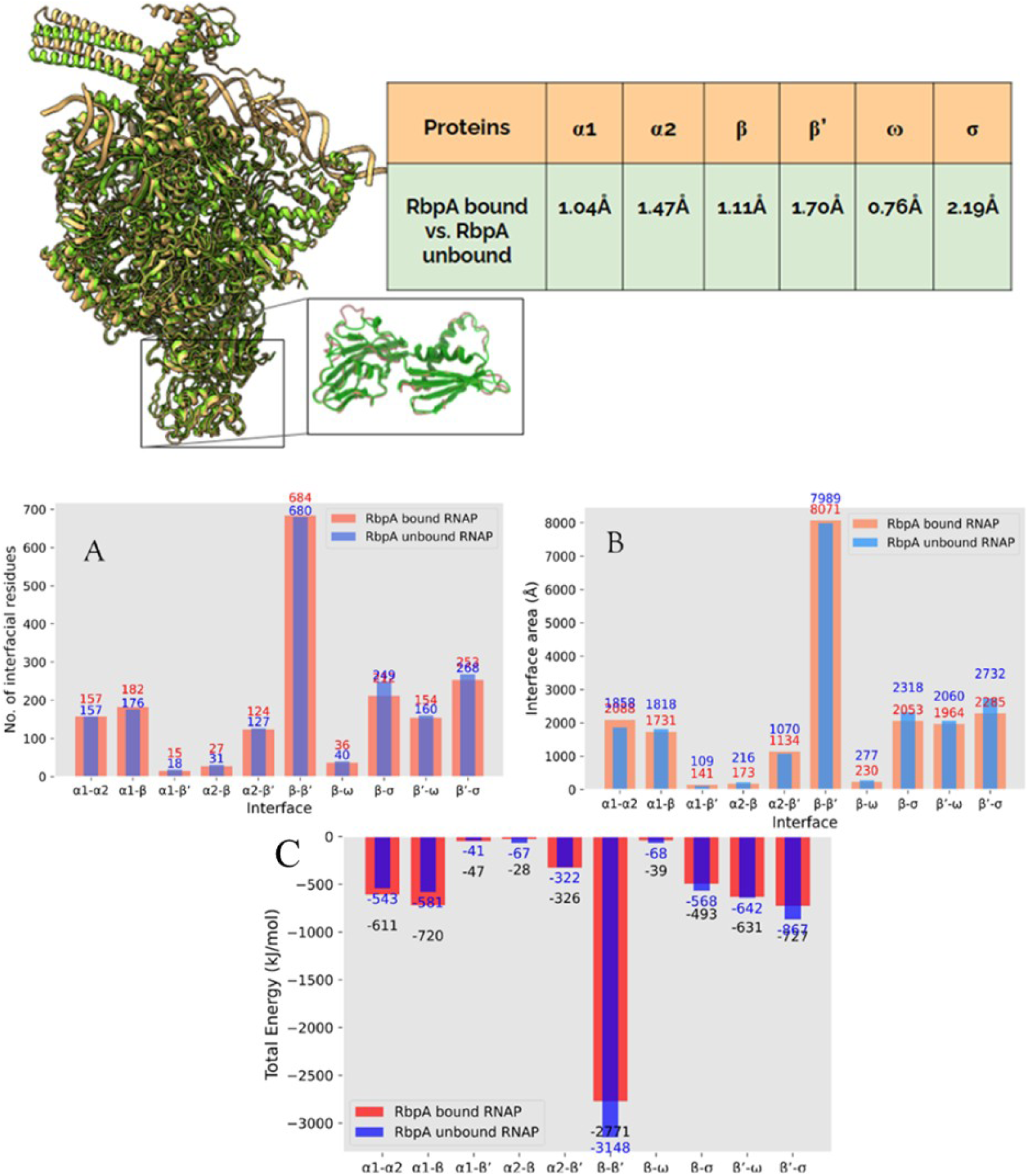
Alignment of RbpA bound and RbpA unbound RNAP holoenzyme complexes. In the inset, aligned *α*1 subunits are shown. The table contains the global RMSD between various subunits. Interfacial properties analysis. Analysis of interfacial properties among various subunits. X-axis is the name of the interface and on the Y-axis, there is the number of interactions or Area of Interaction (Å). A) Total number of interfacial residues, B) Area of interaction C) Number of potential electrostatic interactions, D) Number of salt bridges, E) Number of Van der Waals pairs, and F) Number of hydrophobic interactions.

To gain insight into the binding mechanism of RbpA, we examined the occurrence of various interactions (Electrostatic, Hydrophobic, and salt bridges) among all subunits (Supplementray figure S4B, C, and D), which exhibited consistent trends with the interfacial residue analysis. Moreover, the analysis of the interaction area provided further insights into the mode of interaction. Some subunits demonstrated an increase in the interaction area, while others displayed a decrease. Furthermore, an assessment of the total stabilization energy between subunits indicated that *α*1 and *α*2 exhibited higher levels of stabilization, while the interactions involving other subunits changed to prepare for interaction with DNA (as illustrated in Figure **1**).

Therefore, we can infer that upon interaction with RbpA, subunits *α*1, *α*2 and *β* enhance their interactions by employing electrostatic and hydrophobic forces, while subunits *β*’, *ω* and *σ* adopt an alternative strategy to maintain mobility for subsequent DNA interaction. These findings, in conjunction with the energy analysis, offer valuable insights into the nature and dynamics of the protein-protein interactions within the system. These observations indicate that the interaction of RbpA improves the interactions between *α*1, *α*2, and *β* whereas it decreases the interactions of *β*’, *c*, and *σ*, making them accessible for interaction with incoming DNA. In summary, the investigation of interfacial residues, total stabilization energy, and various interaction parameters has provided important findings regarding the structural changes and interactions within the RNA polymerase complex.

### 3.2 Interaction of RbpA with RNA polymerase changes the hotspot residue network

The hotspot residues of RbpA bound and unbound structures were identified and we learned that the association of RbpA brings changes in the hotspot residue network. Figure 2 highlights the distribution of the hot spot residue network in RbpA bound and unbound forms. The identification of hotspot residues in both RbpA bound and unbound structures allowed us to gain insights into the changes occurring in the hotspot residue network upon the RbpA association. Notably, the association of RbpA was found to induce alterations in the hotspot residue network. By examining the hotspot residues, we could elucidate the crucial regions involved in the protein-protein interactions and their potential functional significance. This analysis of the hotspot residue network enhances our understanding of the molecular mechanisms underlying the interactions between RbpA and its associated proteins.

**FIGURE 2.**
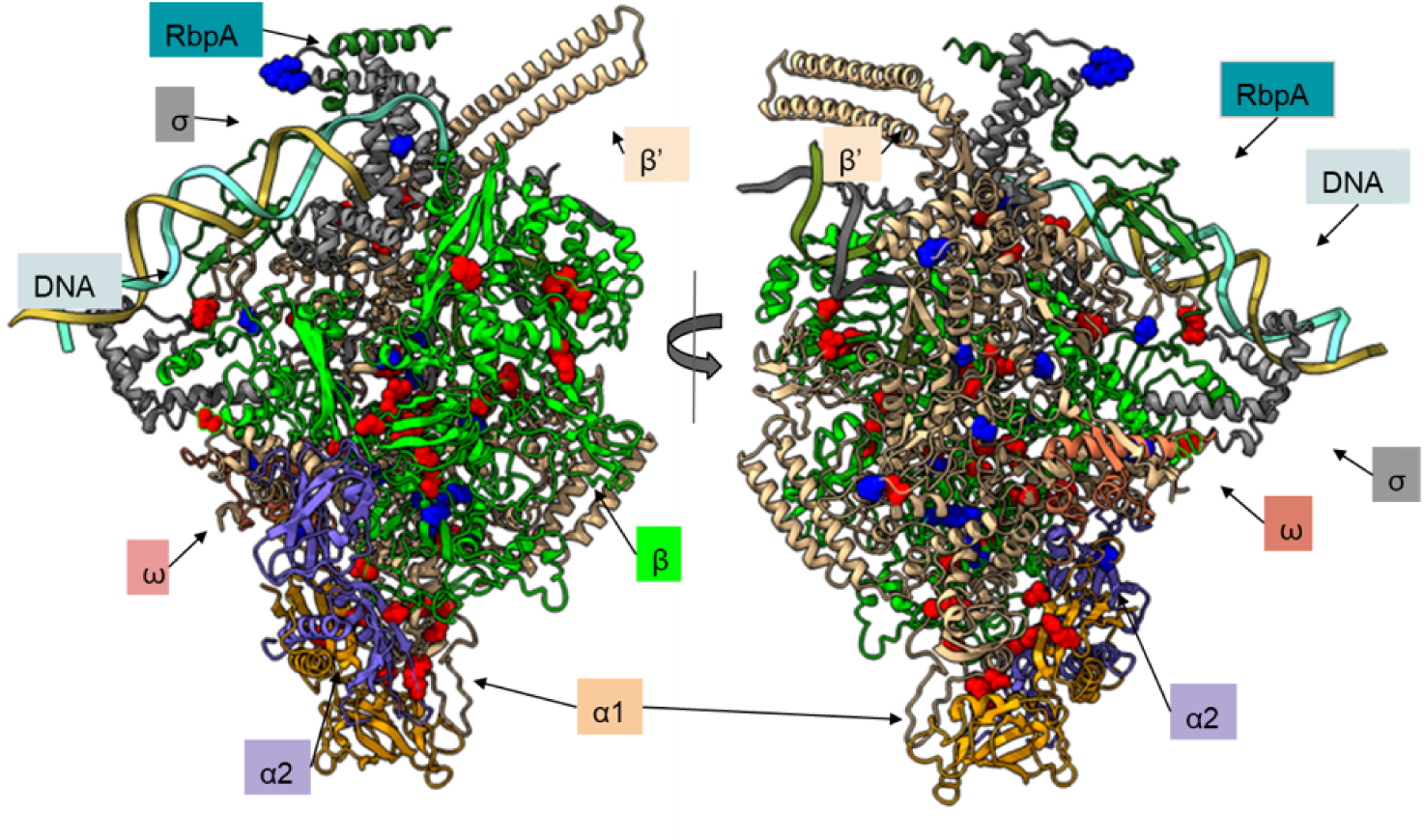
All the hotspot residues of RbpA bound (Red) and unbound (Blue) polymerase complexes were mapped onto its 3D structure of RNA polymerase.

### 3.3 RbpA and CarD interactions alter the dynamics of ***α***1 and ***α***2

*α* subunit of the RNAP complex is composed of 329 amino acid residues. This subunit has three domains, namely N-, C-terminal domains, and a flexible linker domain. It is evident from previous studies that changes in the linker domain can induce alterations in RNAP’s affinity toward DNA promoters [39]. While the*α* subunit primarily serves as a precursor for RNAP assembly, its N- and C-terminal domains perform distinct functions. The N-terminal domains of both *α* subunits form a dimer, acting as a hydrophobic platform facilitating the binding of subunits *β* and *β*’ [40,41].

In our study employing Molecular Dynamics (MD) simulations, we conducted an extensive analysis of the square fluctuations exhibited by all subunits within the different RNA polymerase complexes. We observed that the presence of DNA did not significantly affect the dynamics of either subunit. However, upon binding of RbpA, distinct changes in dynamics were observed in both*α*1 and*α*2 subunits. Specifically, in *α*1, RbpA binding resulted in a shift in dynamics for residues 152-163 (*α*1-*α*2 inter-face) 179-189 (*α*1-*β* interface). Conversely, in *α*2, region 155-165, which corresponds to the interface with *α*1, exhibited increased fluctuations upon interaction with RbpA. These findings are presented through line plots in Figures 3 and 4.

**FIGURE 3.**
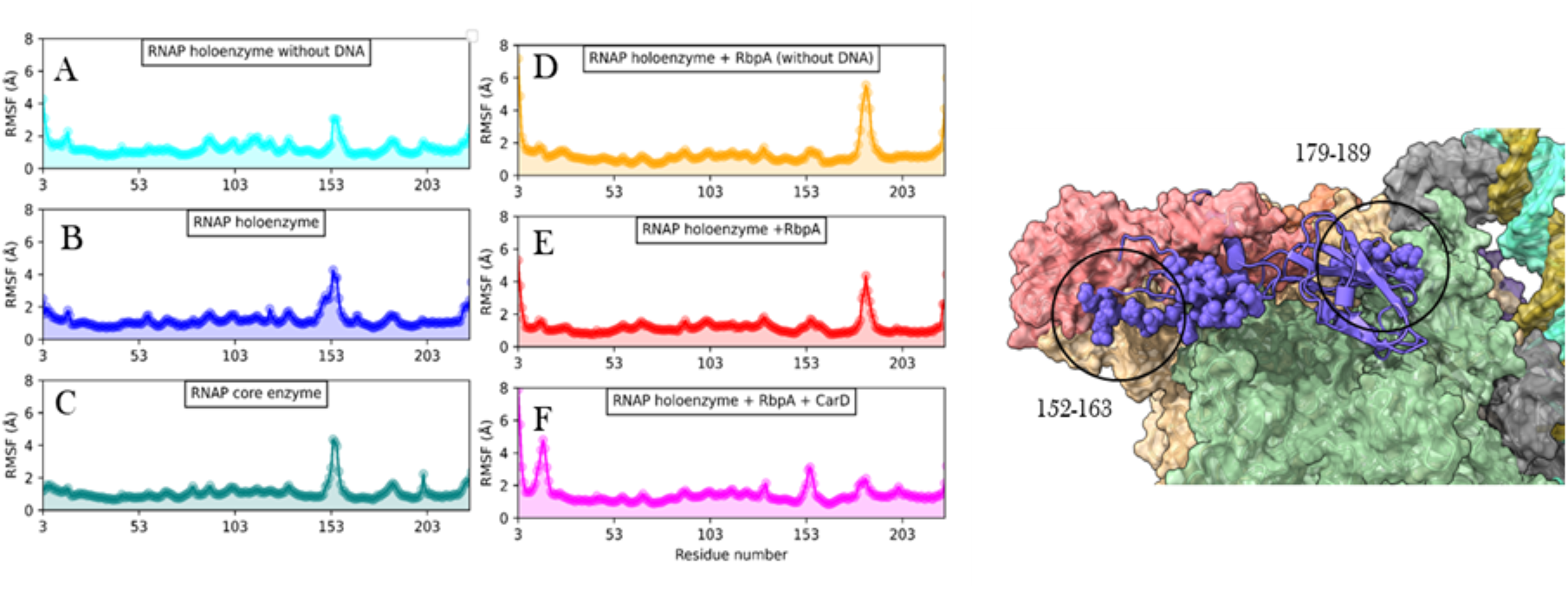
RMSF plots of *α*1 obtained from molecular dynamics analysis. X-axis represents the residue number and Y-axis shows the RMSF in Å. A) RNAP holoenzyme without DNA, B) RNAP holoenzyme, C) RNAP core enzyme, D) RNAP holoenzyme with RbpA but without DNA, E) RNAP holoenzyme with RbpA and DNA, and F) RNAP holoenzyme with RbpA, CarD and DNA. On the right is the visual representation of the residues showing significant differences in the fluctuations. *α*1 is shown in cartoons with highlighted residues as spheres.

**FIGURE 4.**
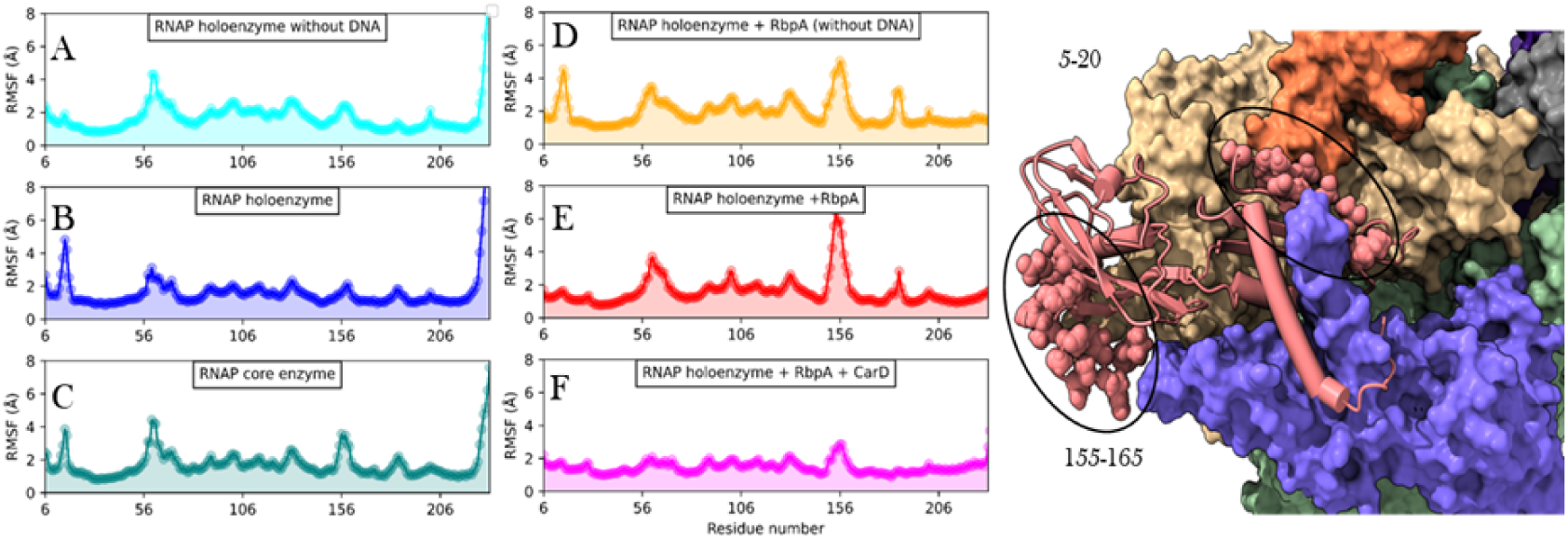
RMSF plots of *α*2 obtained from molecular dynamics analysis. X-axis represents the residue number and Y-axis shows the RMSF in Å. A) RNAP holoenzyme without DNA, B) RNAP holoenzyme, C) RNAP core enzyme, D) RNAP holoenzyme with RbpA but without DNA, E) RNAP holoenzyme with RbpA and DNA, and F) RNAP holoenzyme with RbpA, CarD, and DNA. On the right is the visual representation of the residues showing significant differences in the fluctuations. *α*2 is shown in the cartoon with highlighted residues as spheres.

Furthermore, the binding of CarD in conjunction with RbpA was found to enhance fluctuations in the N-terminal domain *α*1, thereby promoting dynamics at the *α*1-*α*2 interface. However, no significant changes in fluctuations were observed in *α*2 upon the binding of CarD. Additionally, when RbpA and/or CarD were bound to DNA, a decrease in fluctuations was observed at residues 5-20, which corresponds to the interface between *β*’ and and*α*1. Taken together, our findings suggest that RbpA contributes to the stability of the *α*1-*α*2 interface within the RNA polymerase complex, while simultaneously maintaining dynamic interactions with the *β* subunit to facilitate effective DNA interaction. These dynamic interactions and fluctuations are crucial for the proper functioning of the RNA polymerase complex during the process of transcription.

### 3.4 Flexibility of the ***β*** subunit N-terminus is crucial for its interaction with ***β***’ in RNA polymerase

The β’ subunit plays a significant role in transcription, and it encompasses a zinc-binding domain that directly interacts with RbpA and DNA. Within the *β*’ subunit, the *β*’ loop is responsible for regulating DNA movement and the nucleotidyl transfer process. This loop also coordinates with the ‘fork loop’ and ‘bridge helix’ domains to control nucleotide addition during transcription.

Our analysis reveals important insights into the dynamics of the *β*’ loop (Figure 5). We observed that the *β*’ loop displays flexibility in all conformations studied. Interestingly, the binding of RbpA reduces the flexibility of the *β*-*β*’s interface, indicating a stabilizing effect. On the other hand, the presence of CarD reinstates the flexibility of the *β*-*β*’s interface. Notably, the absence of DNA increases the mobility of this region, which may impede the exit of incoming DNA from the transcriptional complex. Furthermore, We found that DNA has a greater impact on the dynamics of the *β*’ subunit compared to the effects induced by RbpA or CarD in the presence of RbpA. These findings suggest that the flexibility of the *β*’ loop is intricately regulated by the presence of RbpA, CarD, and DNA. The ability of the *β*’ loop to modulate DNA movement and coordinate with other regulatory domains underscores its importance in the transcriptional process.

**FIGURE 5.**
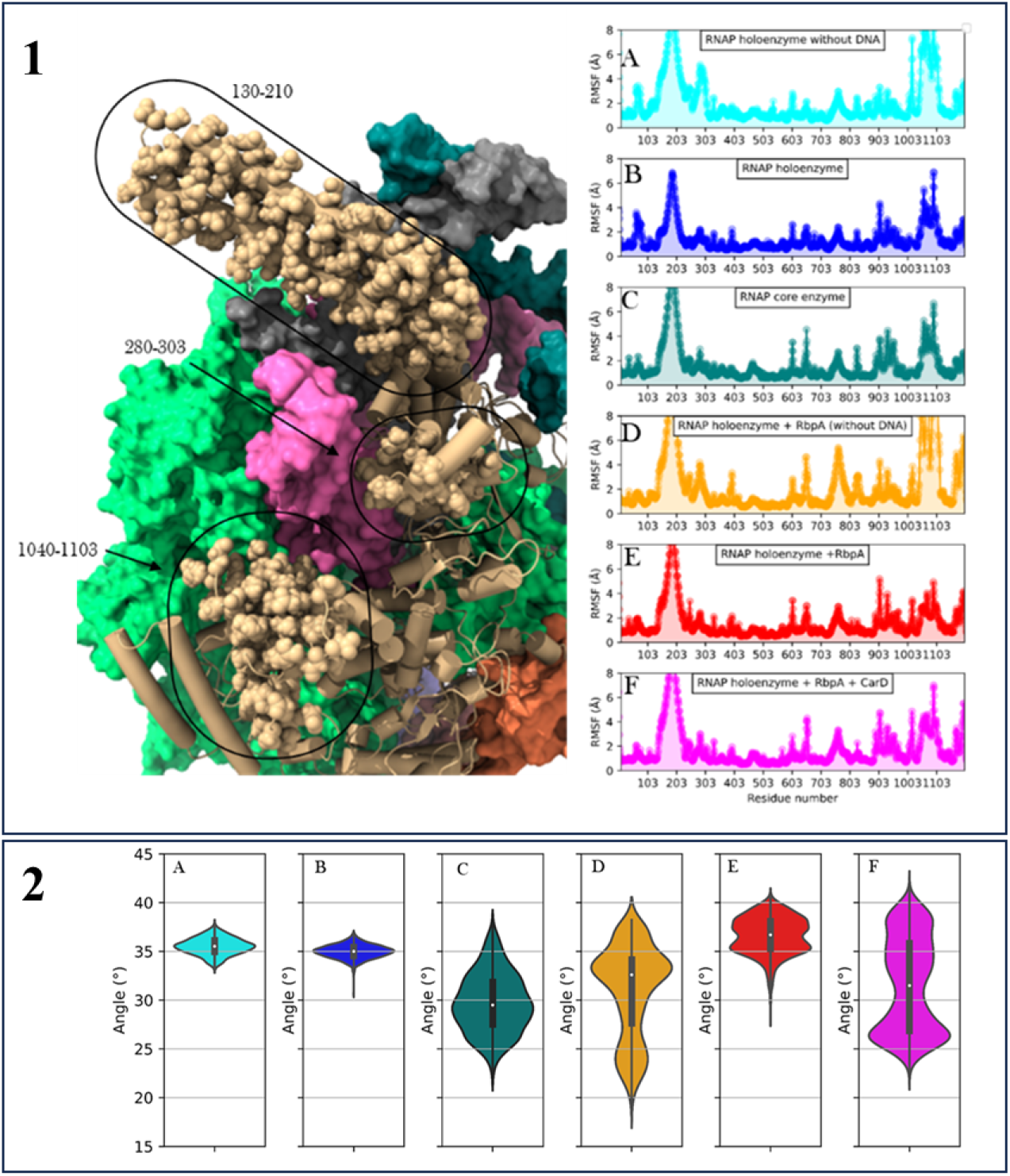
RMSF plots of *β*’ obtained from molecular dynamics analysis. X-axis represents the residue number and Y-axis shows the RMSF in Å. A) RNAP holoenzyme without DNA, B) RNAP holoenzyme, C) RNAP core enzyme, D) RNAP holoenzyme with RbpA but without DNA, E) RNAP holoenzyme with RbpA and DNA, and F) RNAP holoenzyme with RbpA, CarD, and DNA. On the left is the visual representation of the residues showing significant differences in the fluctuations. *β*’ is shown in the cartoon with highlighted residues as spheres. The angle of *β*’ loop with respect to RNAP complex obtained from molecular dynamics analysis. X-axis represents the complexes and Y-axis shows the angle (A) RNAP holoenzyme without DNA, (B) RNAP holoenzyme, (C) RNAP core enzyme, (D) RNAP holoenzyme with RbpA but without DNA, (E) RNAP holoenzyme with RbpA and DNA, and (F) RNAP holoenzyme with RbpA, CarD, and DNA.

### 3.5 RbpA brings in changes in the number of contacts across all the interfaces in the complex

Analysis of inter-protein contacts in RNA polymerase (RNAP) complexes using molecular dynamics (MD) simulations provides insights into the effects of transcription factors and DNA presence on complex stability. The results (Figure 6) demonstrate specific changes in contact patterns within the complex. The observed stability in *α*1-*α*2 and *α*1-*β* interactions suggests their robust nature during complex formation. Conversely, the presence of RbpA influences *α*1-*β*’ interactions in the absence of DNA, indicating a role for RbpA in modulating these interactions. Furthermore, the increased contacts between core enzyme and holoenzyme in the presence of RbpA highlight the influence of this auxiliary factor on complex stability. The lower contacts observed between *β* and *β*’ in the presence of *σ* suggest potential regulatory roles of *σ* in modulating these interactions. Notably, the interactions between *β*-DNA and *β*’-DNA decrease upon binding of *σ*, RbpA, and CarD, indicating their impact on RNAP-DNA interactions. These findings enhance our understanding of the dynamic interplay between transcription factors, DNA, and RNAP subunits during transcription initiation.

**FIGURE 6.**
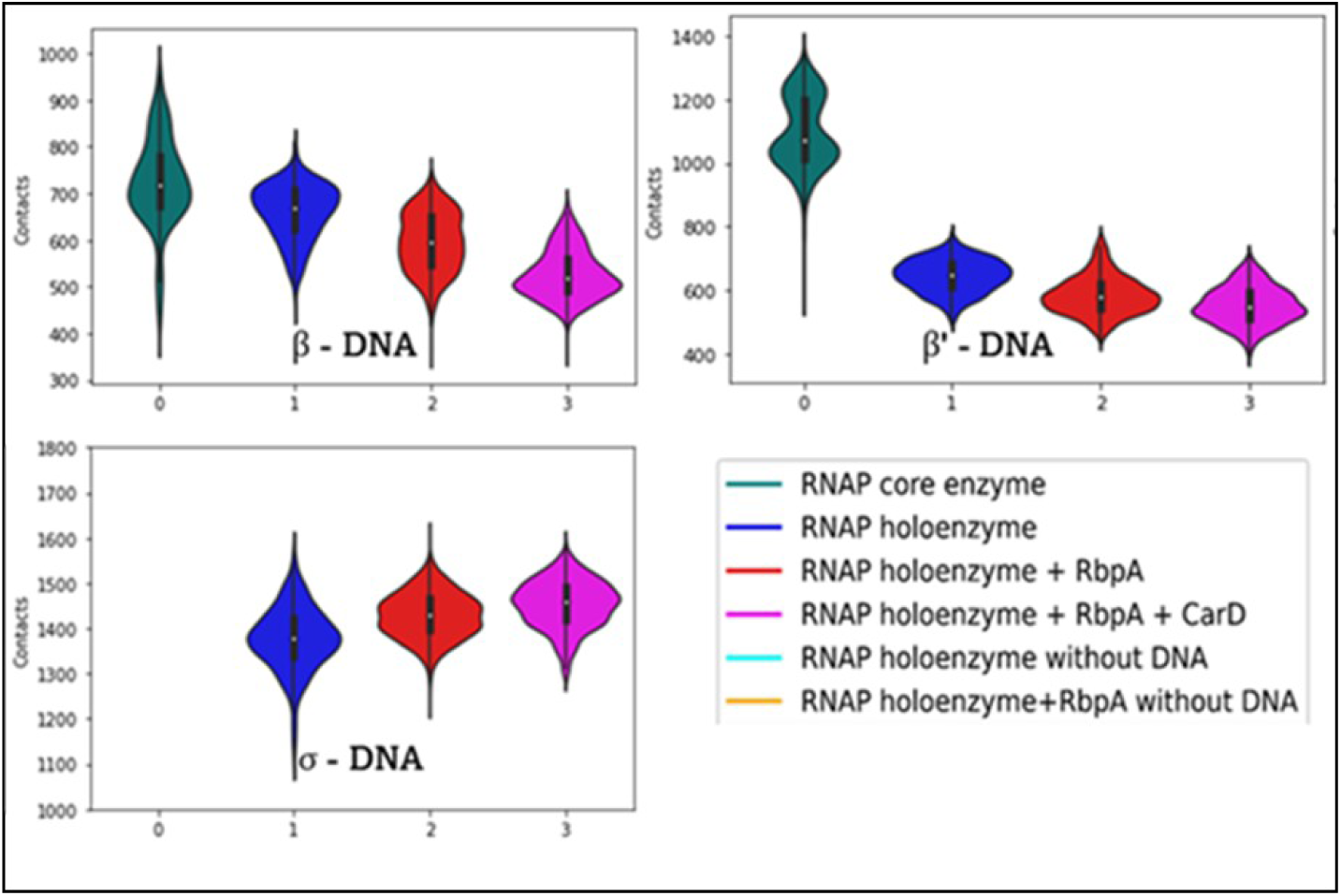
Contact analysis of interfaces from molecular dynamics analysis. Y-axis shows the number of contacts, and the interfaces are marked on the respective graphs and the colors are explained in the figure legend.

### 3.6 Regulation of ***β***’ loop angle by RbpA and CarD

The angle of the *β*’ loop in the RNA polymerase (RNAP) complex is a crucial parameter that influences its function, particularly in regulating the entry of DNA into the complex [42]. In our study, we performed calculations to determine this angle, which represents the orientation of the *β*’ loop relative to the core axis of the *β*’ subunit.

Upon examining our data (Figure **5**), it is evident that there is relatively little variation in the angle of the *β*’ loop between DNA-bound and unbound RNAP holoenzyme complexes. In the absence of the *σ* factor, in the RNAP core enzyme, the *β*’ loop angle remains consistently at approximately 30 degrees. Notably, a significant difference emerges in the distribution of *β*’ loop angles when comparing DNA-bound and unbound complexes in the presence of RbpA. For DNA-unbound complexes, the angle spans a range of 15 to 40 degrees, with an average angle of approximately 33 degrees. Conversely, in DNA-bound RNAP complexes with RbpA, the angle distribution narrows and ranges from 28 to 41 degrees. Notably, there are two distinct peaks on each side of the average angle, with the average itself measuring around 37 degrees. Moreover, the interaction of CarD along with RbpA further influences the *β*’ loop angle. In this case, the angle distribution spans a wider spectrum, ranging from 20 to 42 degrees, with an average angle of approximately 32 degrees. Additionally, a peak was observed at 26 degrees in this distribution. The presence of DNA alone does not significantly alter the angle of the *β*’ loop, indicating that DNA binding alone may not directly influence its conformational changes. However, the presence of RbpA in DNA-bound complexes narrows the distribution of *β*’ loop angles, suggesting a stabilizing effect on the *β*’ loop orientation. This implies that RbpA may contribute to the precise positioning of the *β*’ loop during DNA binding. Furthermore, the interaction of CarD with RbpA expands the range of *β*’ loop angles, potentially indicating a dynamic interplay between CarD, RbpA, and the *β*’ loop, which may be relevant for regulating DNA entry into the RNAP complex.

### 3.7 Comparative Energetic Analysis of RNA Polymerase Complexes

Our mmPBSA analysis revealed distinct energy profiles among the investigated complexes (Figure 7). The total energy values, which reflect the overall stability of the complexes, exhibited a hierarchical order with the holoenzyme bound with RbpA and CarD displaying the highest stability, followed by the holoenzyme with RbpA, holoenzyme alone, and the core enzyme alone. These findings indicate that the presence of RbpA and CarD significantly contribute to the enhanced stability of the holoenzyme. Solvation energy, which characterizes the interaction between the complexes and the surrounding solvent, showed negative values for all complexes, suggesting favorable solvation. Notably, the holoenzyme displayed the highest solvation energy, followed by the holoenzyme with RbpA and CarD, the holoenzyme with RbpA, and the core enzyme. These results indicate that the presence of RbpA and CarD promotes stronger hydration, potentially contributing to the improved stability and functionality of the holoenzyme. Analysis of electrostatic energy, representing the attractive or repulsive forces between charged particles within the complexes, revealed negative values for all complexes, indicating predominantly attractive interactions. The holoenzyme with RbpA exhibited the highest electrostatic energy, followed by the holoenzyme with CarD, the holoenzyme alone, and the core enzyme. These observations suggest that the holoenzyme complexes possess stronger attractive forces between charged particles, further enhancing their stability and functional properties. Van der Waals energy, indicative of the attractive forces between non-polar molecules or regions, showed negative values for all complexes. The holoenzyme with RbpA and holoenzyme with CarD exhibited similar van der Waals energies, suggesting comparable non-polar interactions. The holoenzyme alone and the core enzyme displayed relatively weaker van der Waals forces. These results imply that the presence of RbpA and CarD reinforces favorable non-polar interactions within the holoenzyme. Our observations suggest that the holoenzyme, particularly in the presence of RbpA and CarD, exhibits enhanced stability and improved solvation properties. The stronger electrostatic and van der Waals interactions observed in the holoenzyme complexes further contribute to their stability. These energetic advantages likely contribute to the improved enzymatic activity and functional performance of the holoenzyme.

**FIGURE 7.**
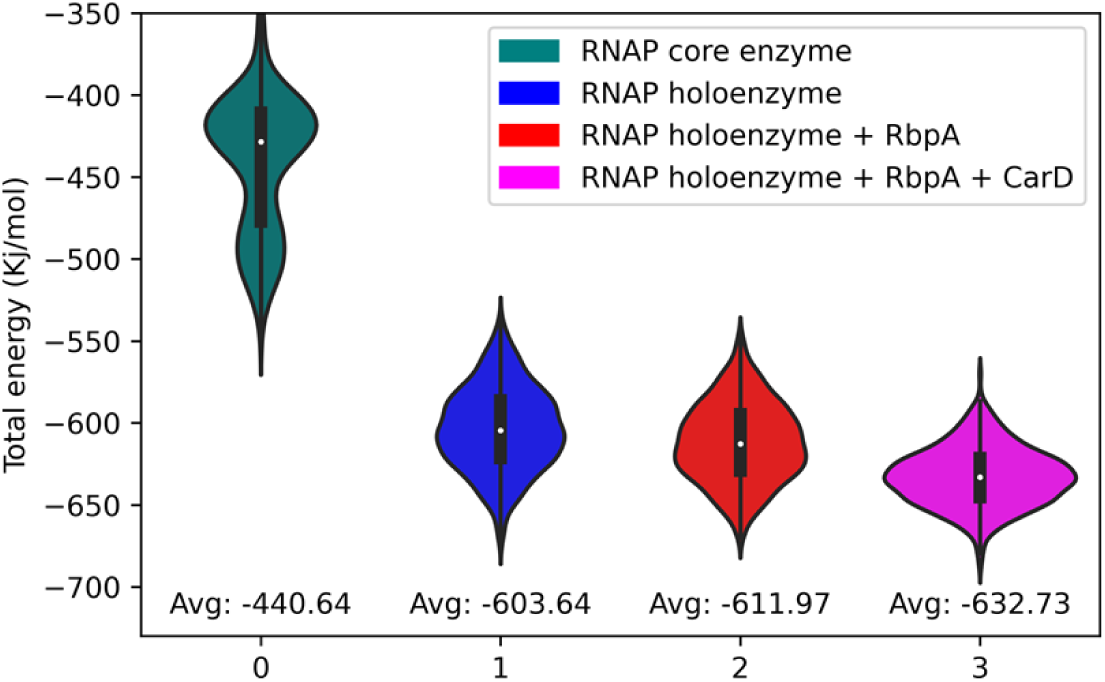
Analysis of interaction energy between complex and DNA obtained from molecular dynamics analysis. X-axis represents the complexes and Y-axis shows the energy in Kj/mol. A) Total energy, B) Solvation energy C) Electrostatic energy and D) Van der Waals energy.

### 3.8 RbpA and CarD interaction with holoenzyme bring changes in the communication among subunits

Analysis of the effectiveness profiles (as described in materials and methods) (Figure 8) revealed distinct patterns upon RbpA binding. The binding of RbpA was found to have minimal impact on the effectiveness profiles of the RNAP holoenzyme sub-units. Conversely, the presence of CarD along with RbpA resulted in improved effectiveness profiles across all subunits, with a notable enhancement observed in the *β* and *β*’ subunits. These findings suggest that CarD plays a pivotal role in modulating the effectiveness and functional dynamics of RNAP holoenzyme, particularly in regions associated with the *β* and *β*’ subunits.

**FIGURE 8.**
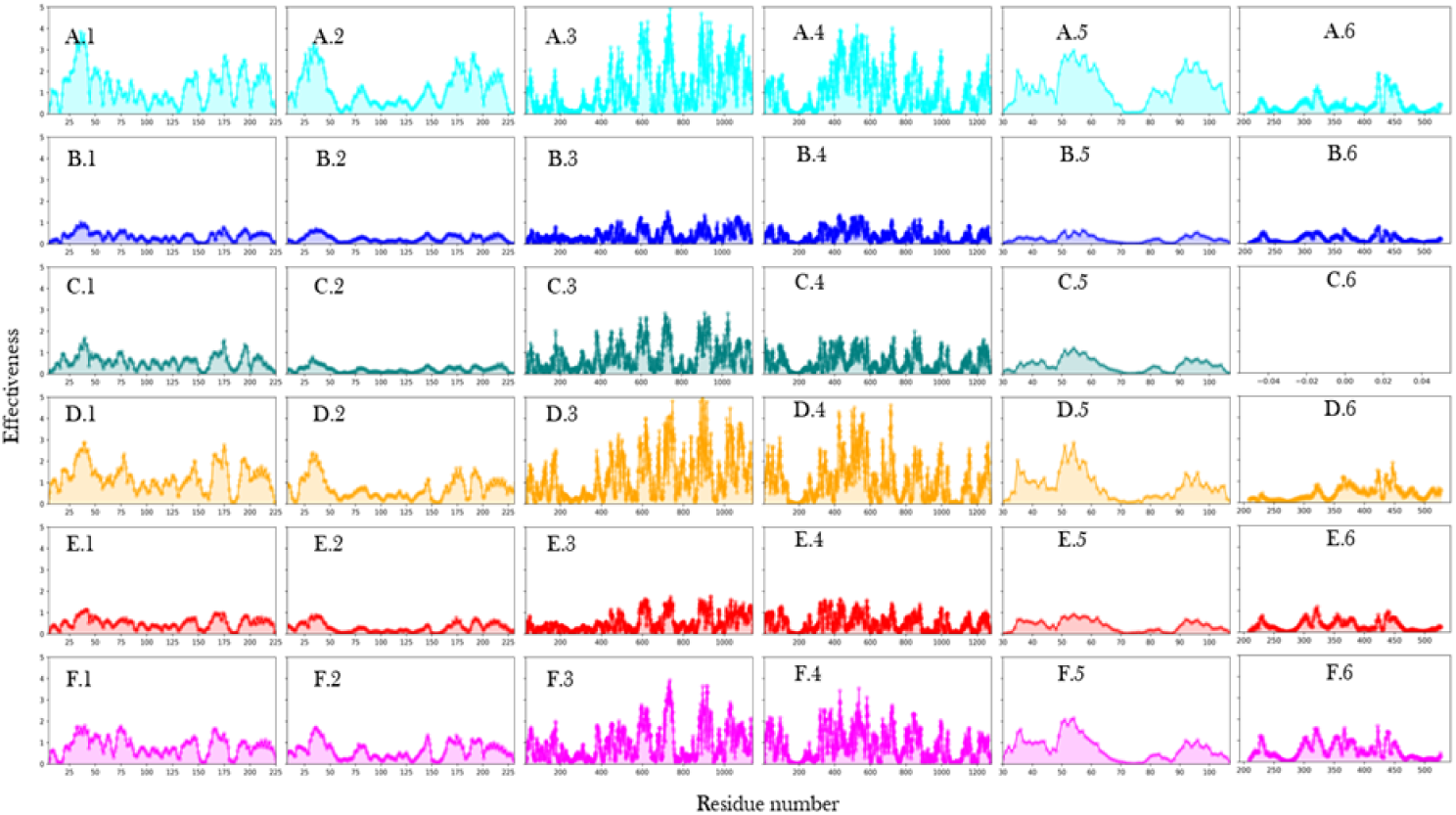
Analysis of effectiveness profiles of various subunits from molecular dynamics analysis. X-axis represents the residues and Y-axis shows the effectiveness. A) RNAP holoenzyme without DNA, B) RNAP holoenzyme, C) RNAP core enzyme, D) RNAP holoenzyme with RbpA but without DNA, E) RNAP holoenzyme with RbpA and DNA, and F) RNAP holoenzyme with RbpA, CarD, and DNA. And 1) *α*1, 2) *α*2, 3) *β*, 4) *β*’, 5) ω and 6) *σ*.

In contrast, the binding of the sigma factor led to a decrease in effectiveness across all subunits of the RNAP holoenzyme. This decrease in effectiveness indicates a dampened response of the RNAP subunits to external stimuli upon sigma factor binding. These results suggest that the sigma factor binding might stabilize the RNAP holoenzyme and reduce its overall effectiveness to ensure specific and regulated transcription initiation. Moreover, the absence of DNA in both the RNA polymerase holoenzyme and its complex with RbpA resulted in increased effectiveness profiles across all residues, particularly in the *β* and *β*’ subunits. These findings indicate that the presence of DNA helps to modulate the sensitivity of the RNAP holoenzyme and its interaction with RbpA, highlighting the importance of DNA in influencing the functional dynamics of the holoenzyme and its associated factors.

Analysis of the sensitivity profiles (Figure 9) demonstrated distinct changes induced by transcription factor binding and the essence of DNA. The binding of RbpA was found to alter the effector residues from positions 150-160 to positions 175-185 *α*1. Subsequent interaction with CarD resulted in the N-terminal residues acting as sensors, indicating a shift in the regulatory mechanisms modulating RNAP holoenzyme dynamics.

**FIGURE 9.**
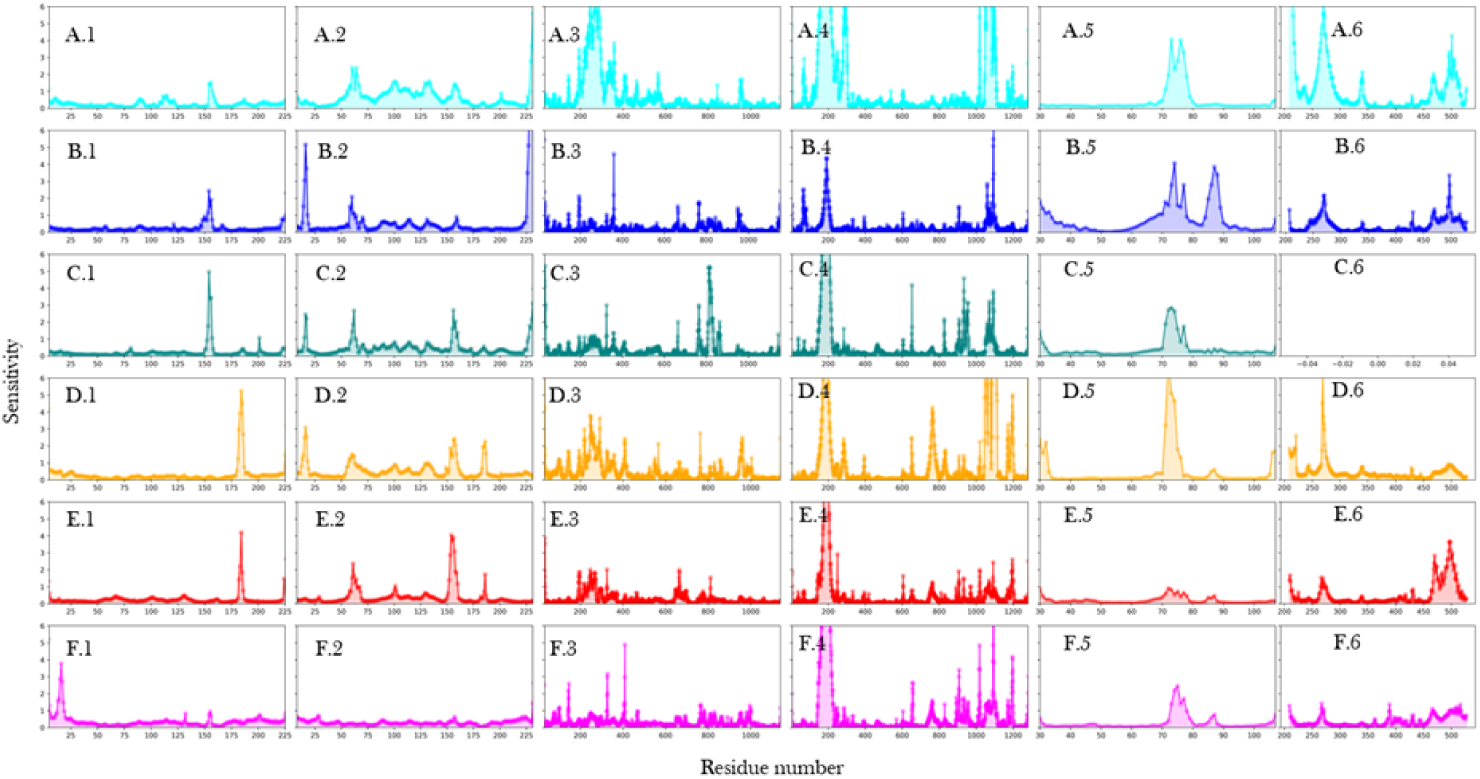
Analysis of sensitivity profiles of various subunits from molecular dynamics analysis. X-axis represents the residues and Y-axis shows sensitivity. A) RNAP holoenzyme without DNA, B) RNAP holoenzyme, C) RNAP core enzyme, D) RNAP holoenzyme with RbpA but without DNA, E) RNAP holoenzyme with RbpA and DNA, and F) RNAP holoenzyme with RbpA, CarD, and DNA. And 1) *α*1, 2) *α*2, 3) *β*, 4) *β*’, 5) ω and 6) *σ*.

In contrast, sigma factor binding decreased sensitivity in the region of residues 150-160 in*α*2, suggesting a dampening effect on the regulatory capacity of these residues. Conversely, RbpA binding upregulated the sensitivity of residues in this region, indicating a potential role for RbpA in enhancing the regulatory capacity of the RNAP holoenzyme. The presence of DNA was found to decrease the sensitivity of residues within the *β* subunit, specifically in the regions of residues 200-400 and 1000-1200. This reduction in sensitivity suggests that DNA binding to the holoenzyme influences the regulatory potential of these residues, potentially affecting the transcription initiation process.

Furthermore, the binding of RbpA and CarD decreased the sensitivity of residues within the *c* subunit in the region of residues 70-80. This finding suggests that the presence of RbpA and CarD may modulate the regulatory role of the *c* subunit in the RNAP holoenzyme. Interestingly, residues 80-90 were identified as sensors in the presence of the *σ* factor and DNA. This observation suggests that the interplay between the *σ* factor and DNA influences the regulatory capacity of these residues, potentially contributing to transcription initiation processes. Additionally, in the absence of DNA, the N-terminal region of the *σ* subunit acted as the sensor. This finding implies that in the absence of DNA, the N-terminal region of *σ* may play a more prominent role in modulating the RNAP holoenzyme dynamics.

### 3.9 Natural ligands interact with the dimer interface of ***α***1 subunit

Virtual screening was performed [43–48] focusing on the α subunits and against two databases of small molecules (as described in Methods). Two potential binding sites were predicted using the sitemap tool of Schrodinger. One of the binding sites (Site 1) is at the interface of α subunits dimer and has site score of 0.894 and DScore of 1.004. Another binding site (site 2) includes residues from α dimer and beta subunit and has site score of 0.929 and DScore 1.032. We used the molecules from both the databases for docking at both site1 and site2.

At site 1, for the HTVS (High-Throughput virtual screening), we retained 10000 ligands, and it resulted in the top hit with a docking score of −7.567. All of the HTVS hits were subjected to SP (Standard precision) docking and “039-337-166” with a docking score of −9.238 was found to be the top hit compound. The top 1000 ligands from SP results were subjected to XP (Extra precision) docking and the best ligand is “002-533-992” with a docking score of −13.257 kcal/mol. All of the XP ligands were re-ranked based on the binding energy calculations from MM-GBSA where limited protein flexibility was considered. “042-624-487” was the top hit with ΔG of −83.5 kcal/mol. The top 10 hits were examined further for their interactions and clashes. The top 10 compounds from each dataset at both sites were subjected to 100ns MD simulations to study the stability of their interactions.

The top compound at site 1 obtained from the molecular dynamics analysis using compounds from Molprot database is 039-339-145 (saikosaponin-F). The complexes were subjected to 100ns MD simulations, and the results are presented in Figure 10. In compound 039-339-145, the majority of the interactions are H-bonding and hydrophobic. Hydrogen bond interactions are from Arg and Leu whereas Phe is a major contributor for hydrophobic interactions. We also observe that the ligand is in contact with the protein throughout the simulation indicating the stability of the complex. The top compound for site4 is discussed in detail in the supplementary information.

**FIGURE 10.**
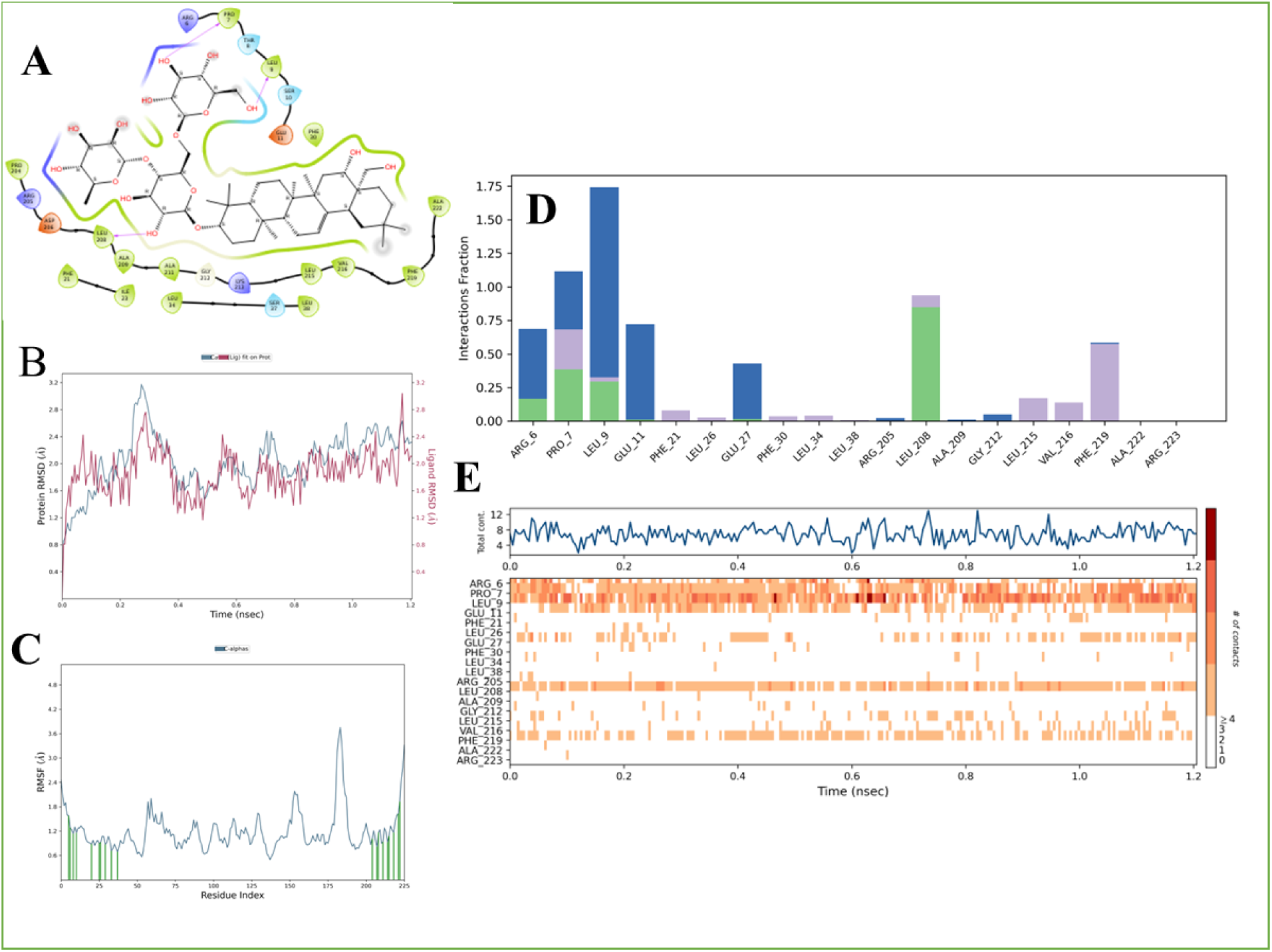
Analysis of the MD simulation results of the compound 039-339-145(saikosaponin-F) with α-subunit of RNA polymerase. A) Pictorial representation of the binding site of the compound 039-339-145 (saikosaponin-F). B) RMSD (Root mean square deviation) of the protein and ligand during the simulation the X-axis contains the time and the Y-axis represents the RMSD. C) RMSF (Root mean square fluctuations) of the residues in α subunit during the simulation. Time is on the X-axis and the Y-axis represents the RMSF. D) Fraction of the interactions of a given residue during the simulation. E) Plot representing the residue involved in the interaction (on the Y-axis) with ligand with simulation time on the X-axis.

### 3.10 FDA approved compounds also interact with ***α***1 at interface of *α* dimer

Among the FDA-approved compounds, Elbasvir is found to be the best compound that interacts with α interface at site1. It is used as a part of combination therapy to treat chronic hepatitis C and was also predicted to bind to RNA-dependent RNA polymerase of SARS-CoV2 [49]. Elbasivir also maintains contact with α subunit throughout the simulation time of 100ns. We also observed that Elbasivir mainly interacts via hydrogen bonding and through the residues from the N-and C-terminal. Residues Met, Ile, and Ala contribute to hydrogen bonding (Figure 11). We followed the above protocol for site 2 which involves α dimer and beta subunit. Likewise, Icotinib was found to stably interact in the site throughout the simulation of 100ns (please see Supplementary information for details).

**FIGURE 11.**
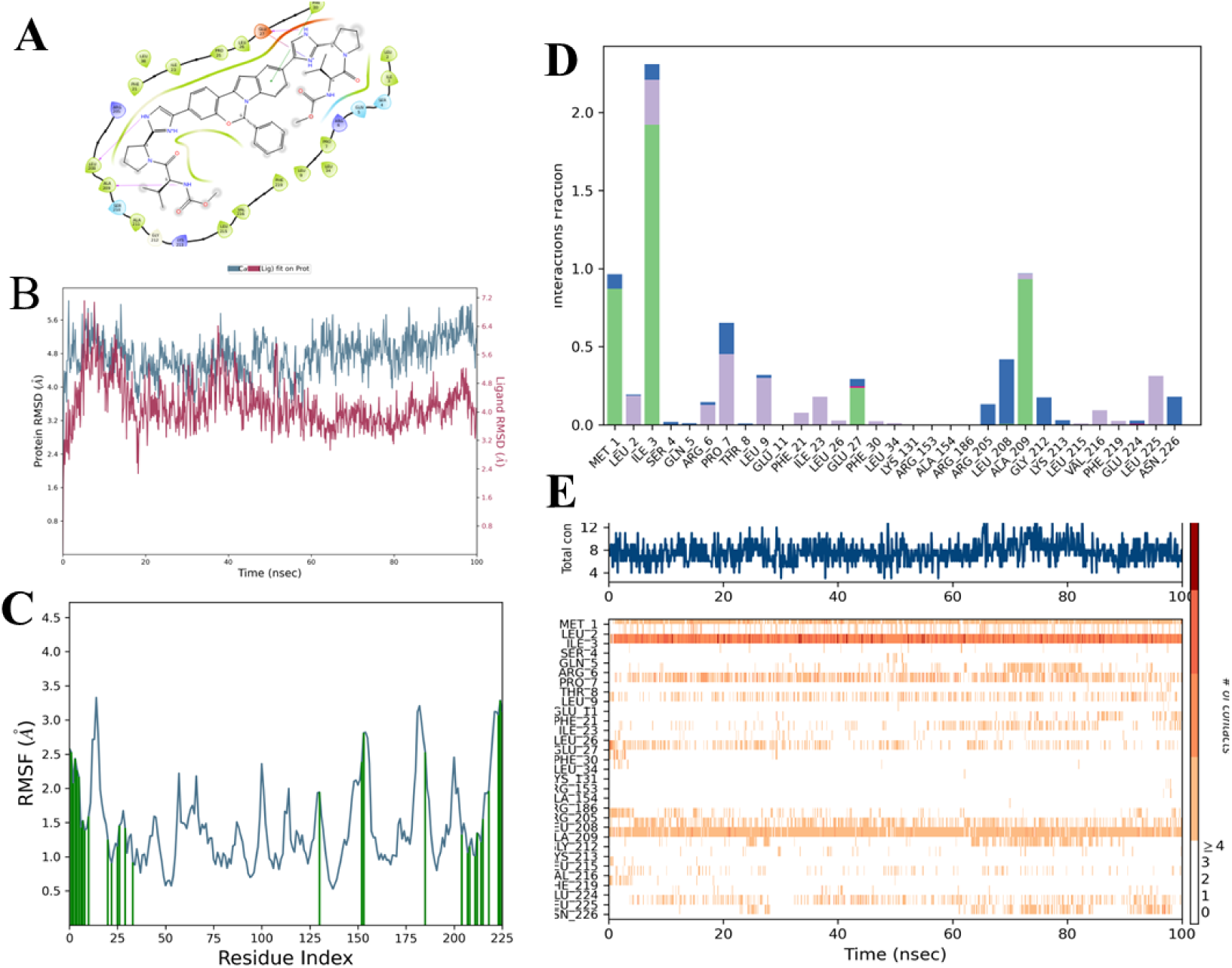
Analysis of the MD simulation results of the compound Elbasivir with α-subunit of RNA polymerase. A) Pictorial representation of the binding site of the compound Elbasivir B) RMSD (Root mean square deviation) of the protein and ligand during the simulation the X-axis contains the time and the Y-axis represents the RMSD. C) RMSF (Root mean square fluctuations) of the residues in α subunit during the simulation. Time is on the X-axis and the Y-axis represents the RMSF. D) Fraction of the interactions of a given residue during the simulation. E) Plot representing the residue involved in the interaction (on the Y-axis) with ligand with simulation time on the X-axis.

## 4 DISCUSSION

This study aimed to unravel the molecular mechanisms underlying the functional regulation of the RbpA protein upon its interaction with RNA polymerase (RNAP). A comprehensive analysis of structural and dynamic changes was conducted in the RNAP complex in the presence of RbpA, DNA, and CarD using molecular dynamics simulations and normal mode analysis. The structural analysis revealed significant differences between RbpA-bound and unbound RNAP complexes, with notable local structural changes observed in the *β*, *β*’, and *σ* subunits. Interfacial interactions and energy analysis provided insights into the nature and dynamics of protein-protein interactions within the system. The findings implied that RbpA enhances interactions between *α*1, *α*2, and *β* subunits while decreasing interactions involving *β*’, *c*, and *σ* subunits, thus facilitating their accessibility for DNA interaction. It could be conceptualized that the *α*1,2 and *β*-*β*’ are held together by the subunit acting like a glue. However, the effectiveness of function is acquired upon binding of RbpA, where the protein subunits rearrange themselves by providing sufficient entropic loss to the system. The fact that the hotspots at *α*1, *α*2 change upon their binding could suggest that this region acts like a “wedge” by accommodating structural changes to the entire complex.

The hotspot network analysis suggested the possibility of allostery and highlighted the involvement of changes in the structure and dynamics of subunits. We employed normal mode analysis to examine global motions in the RNAP complex and observed that RbpA primarily affects the dynamics of the *β*’ loop (Detailed analysis is presented in supporting information). Although normal mode analysis gives us information about 80% of the modes of a protein, we could not observe significant changes in fluctuations in other subunits. So, we performed an all-atom molecular dynamics analysis for 100ns. Molecular dynamics analysis further revealed changes in subunit dynamics in the presence of RbpA, DNA, and CarD. The binding of CarD in conjunction with RbpA increased fluctuations in the N-terminal domain of *α*1, promoting dynamics at the *α*1-*α*2 interface.

However, no significant changes in fluctuations were observed in *α*2 upon the binding of CarD. Additionally, the presence of RbpA and/or CarD, along with DNA, decreased fluctuations at the *β*’-*α*1 interface, indicating their role in stabilizing the complex. The *β*’ loop, crucial for DNA movement and coordination with other regulatory domains, was found to be intricately regulated by RbpA, CarD, and DNA. Furthermore, the study highlighted the role of RbpA in modulating the flexibility of *σ* factor residues involved in functional interactions with RNAP core and DNA during transcription initiation. The holoenzyme complexes, particularly in the presence of RbpA and CarD, exhibited enhanced stability, improved solvation properties, and stronger electrostatic and van der Waals interactions, which likely contributed to their improved enzymatic activity and functional performance. Moreover, the interactions between *β*-DNA and *β*’-DNA were found to decrease upon the binding of *σ*, RbpA, and CarD, suggesting their impact on RNAP-DNA interactions. The absence of DNA revealed the N-terminal region of the *σ* subunit as a prominent sensor, emphasizing its role in modulating RNAP holoenzyme dynamics. The presence of DNA was crucial for modulating the sensitivity of the RNAP holoenzyme and its interaction with RbpA, highlighting the importance of DNA influencing functional dynamics.

Overall, this study provides valuable insights into the structural changes, protein-protein interactions, and dynamics within the RNA polymerase complex upon the binding of RbpA, DNA, and CarD. These findings contribute to the understanding of the dynamic interplay between transcription factors, DNA, and RNAP subunits during transcription initiation, shedding light on the regulatory mechanisms underlying gene expression.

## AUTHOR CONTRIBUTIONS

NS: conceptualization; NS, RS: guidance on analysis; SB: coding, analyses, and first draft of the manuscript; RS: improvement of the manuscript

## DEDICATION

This article is dedicated to one of the authors, late Prof. N. Srinivasan.

## ACKNOWLEDGEMENTS

We thank Indian Institute of Science and NCBS for infrastructural facilities. We thank Dr. Meenakshi Iyer for useful discussions and help in virtual screening. RS is a J.C. Bose National Fellow (JBR/2021/000006) from the Science and Engineering Research Board, India. RS would also like to thank Bioinformatics Centre Grant funded by the Department of Biotechnology, India (BT/PR40187/BTIS/137/9/2021) and the Institute of Bioinformatics and Applied Biotechnology for the funding through her Mazumdar-Shaw Chair in Computational Biology (IBAB/MSCB/182/2022). SB is a CSIR Senior Research fellow and would like to thank CSIR for fellowship support.

## SUPPORTING INFORMATION

Supporting information is provided in the document named ‘Supplementary Information’.

## 4.1 Bibliography References

1. Govindarajan S, Amster-Choder O. Where are things inside a bacterial cell?. 2016

2. Rudner DZ, Losick R. Protein subcellular localization in bacteria.. 2010

3. Allison C, Hughes C. Bacterial swarming: an example of prokaryotic differentiation and multicellular behaviour. Tech. Rep. 4, 1933.

4. Brötz-Oesterhelt H, Bandow JE, Labischinski H. Bacterial proteomics and its role in antibacterial drug discovery. Mass Spectrometry Reviews 2005; 24(4): 549–565. doi: 10.1002/mas.20030

5. Berendsen HJ, Hayward S. Collective protein dynamics in relation to function. 2000

6. Dobson CM. Biophysical Techniques in Structural Biology. Annual Review of Biochemistry 2019; 88(1): 25–33. doi: 10.1146/annurev-biochem-013118-111947

7. Papageorgiou AC, Poudel N, Mattsson J. Protein structure analysis and validation with X-ray crystallography. In: 2178. Humana Press Inc. 2021 (pp. 377–404)

8. Levy D, Chami M, Rigaud JL. Two-dimensional crystallization of membrane proteins: The lipid layer strategy. In: 504. No longer published by Elsevier; 2001: 187–193

9. Rammohan J, Manzano AR, Garner AL, Stallings CL, Galburt EA. CarD stabilizes mycobacterial open complexes via a two-tiered kinetic mechanism. Nucleic Acids Research 2015; 43(6): 3272–3285. doi: 10.1093/NAR/GKV078

10. Srivastava DB, Leon K, Osmundson J, et al. Structure and function of CarD, an essential mycobacterial transcription factor. Proceedings of the National Academy of Sciences of the United States of America 2013; 110(31): 12619–12624. doi: 10.1073/PNAS.1308270110

11. Stallings CL, Stephanou NC, Chu L, Hochschild A, Nickels BE, Glickman MS. CarD Is an Essential Regulator of rRNA Transcription Required for Mycobacterium tuberculosis Persistence. Journal of End-to-End-testing 2009; 138(1): 146–159. doi: 10.1016/S9999-9994(09)20365-5

12. Dey A, Verma A, Microbiology DC, 2010 u. Role of an RNA polymerase interacting protein, MsRbpA, from Mycobacterium smegmatis in phenotypic tolerance to rifampicin. microbiologyresearch.org 2010; 156(3): 873–883. doi: 10.1099/mic.0.033670-0

13. Hu Y, Morichaud Z, Chen S, Leonetti JP, Brodolin K. Mycobacterium tuberculosis RbpA protein is a new type of transcriptional activator that stabilizes the *σ* a-containing RNA polymerase holoenzyme. Nucleic Acids Research 2012; 40(14): 6547–6557. doi: 10.1093/NAR/GKS346

14. Hubin EA, Tabib-Salazar A, Humphrey LJ, et al. Structural, functional, and genetic analyses of the actinobacterial transcription factor RbpA. Proceedings of the National Academy of Sciences of the United States of America 2015; 112(23): 7171–7176. doi: 10.1073/PNAS.1504942112

15. Boyaci H, Chen J, Lilic M, et al. Fidaxomicin jams mycobacterium tuberculosis RNA polymerase motions needed for initiation via RBPA contacts. eLife 2018; 7. doi: 10.7554/eLife.34823

16. Lin W, Mandal S, Degen D, et al. Structural Basis of Mycobacterium tuberculosis Transcription and Transcription Inhibition. Molecular Cell 2017; 66(2): 169–179. doi: 10.1016/j.molcel.2017.03.001

17. Melo F, Sali A. Fold assessment for comparative protein structure modeling. Wiley Online Library 2007; 16(11): 2412–2426. doi: 10.1110/ps.072895107

18. Zhang Y, Skolnick J. TM-align: A protein structure alignment algorithm based on the TM-score. Nucleic Acids Research 2005; 33(7): 2302–2309. doi: 10.1093/nar/gki524

19. Tina KG, Bhadra R, Srinivasan N. PIC: Protein Interactions Calculator. Nucleic Acids Research 2007; 35(SUPPL.2): 473– 476. doi: 10.1093/nar/gkm423

20. Sukhwal A, Sowdhamini R. PPCheck: A Webserver for the Quantitative Analysis of Protein– Protein Interfaces and Prediction of Residue Hotspots. Bioinformatics and Biology Insights 2015; 9: 141. doi: 10.4137/BBI.S25928

21. Keskin O, Ma B, Nussinov R. Hot regions in protein-protein interactions: The organization and contribution of structurally conserved hot spot residues. Journal of Molecular Biology 2005; 345(5): 1281–1294. doi: 10.1016/j.jmb.2004.10.077

22. Boyaci H, Chen J, Jansen R, Darst SA, Campbell EA. Structures of an RNA polymerase promoter melting intermediate elucidate DNA unwinding. Nature 2019; 565(7739): 382–385. doi: 10.1038/S41586-018-0840-5

23. Gowers RJ, Linke M, Barnoud J, et al. MDAnalysis: A Python Package for the Rapid Analysis of Molecular Dynamics Simulations. PROC. OF THE 15th PYTHON IN SCIENCE CONF 2016.

24. Michaud-Agrawal N, Denning EJ, Woolf TB, Beckstein O. MDAnalysis: A toolkit for the analysis of molecular dynamics simulations. Journal of Computational Chemistry 2011; 32(10): 2319–2327. doi: 10.1002/JCC.21787

25. Bakan A, Meireles LM, Bahar I. ProDy: Protein dynamics inferred from theory and experiments. Bioinformatics 2011; 27(11): 1575–1577. doi: 10.1093/bioinformatics/btr168

26. Banerjee P, Erehman J, Gohlke BO, Wilhelm T, Preissner R, Dunkel M. Super Natural II—a database of natural products. Nucleic Acids Research 2015; 43: D935–D939. doi: 10.1093/NAR/GKU886

27. Wishart DS, Feunang YD, Guo AC, et al. DrugBank 5.0: a major update to the DrugBank database for 2018. Nucleic Acids Research 2018; 46: D1074–D1082. doi: 10.1093/NAR/GKX1037

28. Halgren TA. Identifying and characterizing binding sites and assessing druggability. Journal of Chemical Information and Modeling 2009; 49: 377–389. doi: 10.1021/CI800324M/ASSET/IMAGES/CI-2008-00324M_M006.GIF

29. Halgren T. New Method for Fast and Accurate Binding-site Identification and Analysis. Chemical Biology Drug Design 2007; 69: 146–148. doi: 10.1111/J.1747-0285.2007.00483.X

30. Glide: A New Approach for Rapid, Accurate Docking and Scoring. 1. Method and Assessment of Docking Accuracy. Journal of Medicinal Chemistry 2004; 47: 1739–1749. doi: 10.1021/JM0306430

31. Halgren TA, Murphy RB, Friesner RA, et al. Glide: A New Approach for Rapid, Accurate Docking and Scoring. 2. Enrichment Factors in Database Screening. Journal of Medicinal Chemistry 2004; 47: 1750–1759. doi: 10.1021/JM030644S

32. Friesner RA, Murphy RB, Repasky MP, et al. Extra precision glide: Docking and scoring incorporating a model of hydrophobic enclosure for protein-ligand complexes. Journal of Medicinal Chemistry 2006; 49: 6177–6196. doi: 10.1021/JM051256O

33. Morris GM, Goodsell DS, Halliday RS, et al. Automated Docking Using a Lamarckian Genetic Algorithm and an Empirical Binding Free Energy Function. Journal of Computational Chemistry 1639; 19: 16391662. doi: 10.1002/(SICI)1096-987X(19981115)

34. Bowers KJ, Chow E, Xu H, et al. Scalable algorithms for molecular dynamics simulations on commodity clusters. Proceedings of the 2006 ACM/IEEE Conference on Supercomputing, SC’06 2006. doi: 10.1145/1188455.1188544

35. Sastry GM, Adzhigirey M, Day T, Annabhimoju R, Sherman W. Protein and ligand preparation: Parameters, protocols, and influence on virtual screening enrichments. Journal of Computer-Aided Molecular Design 2013; 27: 221–234. doi: 10.1007/S10822-013-9644-8

36. Harder E, Damm W, Maple J, et al. OPLS3: A Force Field Providing Broad Coverage of Drug-like Small Molecules and Proteins. Journal of Chemical Theory and Computation 2016; 12: 281–296. doi: 10.1021/ACS.JCTC.5B00864

37. Holm L, Kääriäinen S, Rosenström P, Schenkel A. Searching protein structure databases with DaliLite v.3. Bioinformatics 2008; 24(23): 2780–2781. doi: 10.1093/bioinformatics/btn507

38. Gutmanas A, Alhroub Y, Battle GM, et al. PDBe: Protein data bank in Europe. Nucleic Acids Research 2014; 42(D1): D285–D291. doi: 10.1093/nar/gkt1180

39. Meng W. The Escherichia coli RNA polymerase alpha subunit linker: length requirements for transcription activation at CRP-dependent promoters. The EMBO Journal 2000; 19(7): 1555–1566. doi: 10.1093/EMBOJ/19.7.1555

40. Igarashi K, Fujita N, Ishihama A. Identification of a subunit assembly domain in the alpha subunit of Escherichia coli RNA polymerase. Journal of Molecular Biology 1991; 218(1): 1–6. doi: 10.1016/0022-2836(91)90865-4

41. Murakami K, Kimura M, Owens JT, Meares CF, Ishihama A. The two *α* subunits of Escherichia coli RNA polymerase are asymmetrically arranged and contact different halves of the DNA upstream element. Proceedings of the National Academy of Sciences of the United States of America 1997; 94(5): 1709–1714. doi: 10.1073/PNAS.94.5.1709

42. Wang D, Bushnell DA, Westover KD, Kaplan CD, Kornberg RD. Structural Basis of Transcription: Role of the Trigger Loop in Substrate Specificity and Catalysis. Cell 2006; 127(5): 941–954. doi: 10.1016/J.CELL.2006.11.023

43. Gandhimathi, A., & Sowdhamini, R. (2016). Molecular modelling of human 5-hydroxytryptamine receptor (5-HT2A) and virtual screening studies towards the identification of agonist and antagonist molecules. Journal of biomolecular structure & dynamics, 34(5), 952–970. 10.1080/07391102.2015.1062802

44. Sharma, A., Tiwari, V., & Sowdhamini, R. (2020). Computational search for potential COVID-19 drugs from FDAapproved drugs and small molecules of natural origin identifies several anti-virals and plant products. Journal of biosciences, 45(1), 100. 10.1007/s12038-020-00069-8

45. Tiwari V, Viswanath S. Identification of potential modulators of IFITM3 by in-silico modeling and virtual screening. Scientific Reports 2022; 12. doi: 10.1038/S41598-022-20259-8

46. Tiwari, V., & Sowdhamini, R. (2023). Structure modelling of odorant receptor from Aedes aegypti and identification of potential repellent molecules. Computational and structural biotechnology journal, 21, 2204–2214. 10.1016/j.csbj.2023.03.005

47. Abhishek Sharma, Sudhir Krishna, Ramanathan Sowdhamini, Bioinformatics Analysis of Mutations Sheds Light on the Evolution of Dengue NS1 Protein With Implications in the Identification of Potential Functional and Druggable Sites, Molecular Biology and Evolution, Volume 40, Issue 3, March 2023, msad033, 10.1093/molbev/msad033

48. Shailya Verma, Purushotham Reddy and R. Sowdhamini (2023) Integrated approaches for the recognition of small molecule inhibitors for Toll-like receptor 4. Computational and Structural Biotechnology Journal, 21, 3680–3689 10.1016/j.csbj.2023.07.026.

49. Balasubramaniam M, Reis RJS. Computational target-based drug repurposing of elbasvir, an antiviral drug predicted to bind multiple SARS-CoV-2 proteins. ChemRxiv 2020. doi: 10.26434/CHEMRXIV.12084822

